# Two-fluid dynamics and micron-thin boundary layers shape cytoplasmic flows in early *Drosophila* embryos

**DOI:** 10.1101/2023.03.16.532979

**Authors:** Claudio Hernández López, Alberto Puliafito, Yitong Xu, Ziqi Lu, Stefano Di Talia, Massimo Vergassola

**Affiliations:** École Normale Supérieure, 75005 Paris, France; Department of Oncology, University of Turin, 10060 Candiolo, Italy; Candiolo Cancer Institute, FPO - IRCCS, Str. Prov. 142, km 3.95, 10060 Candiolo, Italy; Department of Cell Biology, Duke University Medical Center, Durham, NC 27710 USA; Department of Physics, University of California San Diego, San Diego, CA 92075, USA

## Abstract

Cytoplasmic flows are widely emerging as key functional players in development. In early *Drosophila* embryos, flows drive the spreading of nuclei across the embryo. Here, we combine hydrodynamic modeling with quantitative imaging to develop a two-fluid model that features an active actomyosin gel and a passive viscous cytosol. Gel contractility is controlled by the cell cycle oscillator, the two fluids being coupled by friction. In addition to recapitulating experimental flow patterns, our model explains observations that remained elusive, and makes a series of new predictions. First, the model captures the vorticity of cytosolic flows, which highlights deviations from Stokes’ flow that were observed experimentally but remained unexplained. Second, the model reveals strong differences in the gel and cytosol motion. In particular, a micron-sized boundary layer is predicted close to the cortex, where the gel slides tangentially whilst the cytosolic flow cannot slip. Third, the model unveils a mechanism that stabilizes the spreading of nuclei with respect to perturbations of their initial positions. This self-correcting mechanism is argued to be functionally important for proper nuclear spreading. Fourth, we use our model to analyze the effects of flows on the transport of the morphogen Bicoid, and the establishment of its gradients. Finally, the model predicts that the flow strength should be reduced if the shape of the domain is more round, which is experimentally confirmed in *Drosophila* mutants. Thus, our two-fluid model explains flows and nuclear positioning in early *Drosophila*, while making predictions that suggest novel future experiments.

---

Cytoplasmic flows are ubiquitous in biology, ranging from flows in large *Physarum* cells [1–3] to flows in the extra-cellular space controlling left-right asymmetry in development [4, 5]. Oogenesis and embryogenesis are two biological processes where flows play key roles [6, 7]. Flows are central in both *C. elegans* and *Drosophila* for the specification of oocytes [8–13] and play a crucial role in nuclear and spindle positioning in oocyte mouse meiosis [14, 15]. In most species, cytoplasmic flows are observed in the early stage of embryogenesis [16–19]. While the functional role of these flows is not fully understood, we have recently demonstrated that flows drive proper nuclear positioning in early *Drosophila* [20]. The early fly embryo develops as a syncytium, i.e., a multinucleated cell where molecules are free to diffuse. The embryo is large (about 500*μm* in length) and thus nuclei following fertilization have to migrate distances as large as 200 − 300*μm* to fill the entire embryo [18, 20, 21]. Nuclear movements must be fast, as development proceeds very rapidly to avoid predation and pathogens’ infection [22]. Upon fertilization, nuclei undergo fast and synchronous cycles of cleavage divisions, lasting about 8 minutes each [23, 24]. We have previously shown that during those divisions, flows transport nuclei along the anterior-posterior (AP) axis ensuring that they occupy the entire embryo uniformly [20].

Cytoplasmic flows in early *Drosophila* arise from the coupling between cell cycle oscillations and actomyosin contractility [18, 20, 21]. At mitotic exit, the activity of Cdk1, a master regulator of the cell cycle, begins to decrease near the chromosomes [25–27]. This local down-regulation triggers the activation of the mitotic phosphatase PP1 (and likely PP2A as well) which, together with downregulation of Cdk1, can effectively dephosphorylate mitotic targets in a region of ∼50*μm* around the nuclei [20]. This size is sufficient to trigger dephosphorylation of mitotic targets near the cortex. Thus, PP1 activity effectively couples nuclear and cortical dynamics. Higher PP1 activity at mitotic exit triggers the differential activation of actomyosin contractility at the cortex. Thus, nuclei can drive the spatiotemporal pattern of cortical actomyosin contractility through the regulation of the cell cycle oscillator. Notably, when cortical actomyosin contractility is blocked via optogenetic perturbations, flows are abolished and nuclei do not spread properly [20]. As a result of this reduced nuclear movement, nuclear density across the embryos is highly non-uniform. In turn, this results in asynchronous cell cycles prior to the maternal-to-zygotic transition (MZT), which demonstrates the functional role of the flows [20, 28, 29].

Our previous experimental results strongly argue that cytoplasmic flows observed in early embryos are driven by cortical contractility [20]. Yet, we are still lacking a quantitative picture of how cortical contractions drive the flow of cytoplasm across the embryo and its consequences. Previous models have described the cytoplasm as a viscous fluid with cortical flows imposed as boundary conditions [16, 20, 30]. This model presents severe limitations, both fundamental and practical. First, the model ignores interactions of the actomyosin network at the cortex and the cytosol. In particular, the cytosol should obey the usual no-slip boundary conditions of normal fluids, which is not the case in the above model. Moreover, our analysis of the cytoplasmic vorticity field shows that its small-scale structure in early *Drosophila* embryos deviates from the behavior of a simple viscous fluid. Specifically, vorticity of a Stokes’ flow is a harmonic function and should then have maxima and minima only at the boundary whilst they are experimentally observed in the interior of the embryo [20].

A more advanced physical framework for cytoplasmic flow is offered by models considering the possibility of multiple fluids [31]. However, these models can become very cumbersome with a large number of mathematical terms and couplings. A simpler approach is offered by models inspired by poroelasticity, which were previously shown to capture the dynamics of blebbing and response to microindentation [32–34]. Here, we present a two-fluid physical model for cytoplasmic flows in *Drosophila* embryos. In our model, we consider the interactions between a gel, the contractile actomyosin network, and a passive viscous fluid, the cytosol. We first validate this model by showing that it can capture the basic properties of cytoplasmic flows and then develop its predictions and functional consequences.

## I. RESULTS

### A. Stokes’ flows fail to explain cytoplasmic streaming

To gain insight into the physics of cytoplasmic flows, let us start with the simplest possible option: a single fluid obeying Stokes’ equations, which describe a normal fluid dominated by viscous effects. The Navier-Stokes equations for an incompressible fluid reduce then to [35]:

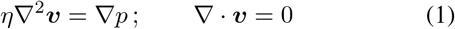

where *η* is the dynamic viscosity, ***v*** the velocity field, and *p* the pressure field.

Three main arguments can be put forward against such a simple description of the cytoplasm. The first two arguments are empirical and data-driven. First, we found that the Stokes’ flows that fit a set of measured data are unable to correctly predict the rest of said data. Specifically, we found that, in order to fit the velocities in the bulk of the embryo, the flow speeds near the embryo cortex (≲ 10*μm*) must rapidly increase and would therefore significantly exceed the measured flow. That suggests that Stokes’ flows cannot capture the behavior of cytoplasmic flows near the embryo boundary. Second, while the large-scale patterns are visually similar, quantitative properties of the experimental flows clearly deviate from Stokes’ flows. A convenient way to highlight those effects is the vorticity field *ω*_*v*_ = ∇ × ***v*** [35]. This quantity measures the local circulation of fluid elements and the presence of a derivative in its definition highlights small-scale properties of the velocity ***v***. From Eq. (1), we can derive:

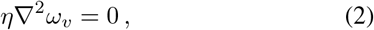

which implies that the vorticity *ω*_*v*_ is a harmonic function. This is relevant as an elementary theorem states that the extrema of harmonic functions must be located at the boundary of the domain. Conversely, the vorticity of our measured flows systematically features four extrema well inside the embryos [20], which manifestly demonstrates deviations of cytoplasmic flows from a Stokes’ structure.

The third argument on boundary conditions relates to the first point above. A normal fluid obeys no-slip conditions at the boundary, i.e., it should move at the same velocity as the boundary. Since we do not observe the vitelline membrane undergoing significant movement, we can assume that the perivitelline fluid should be at rest at the boundary. In the preblastoderm stage, the plasma membrane is in most places adjacent to the vitelline membrane, thus no-slip boundary conditions are a reasonable approximation, which contradicts imposing a non-trivial velocity as boundary condition.

In sum, a single passive Stokes flow is insufficient and a better physical model is needed to capture the dynamics of flows in early *Drosophila* embryos. As we show in the next Section, a parsimonious way out of these limitations is the introduction of two fluids, which will be shown to lead to the formation of a boundary layer near the cortex and differential motion of gel and cytosol.

### B. Two-fluid model for cytoplasmic flows

To establish a more relevant model for cytoplasmic flows, we explicitly consider both the actomyosin network and the cytosol, described as an active gel and a passive viscous fluid, respectively. We describe the interaction between the two fluids with a simple friction term. Moreover, we include the nuclei and their cell cycle regulation to obtain a model that can describe nuclear positioning by the flows. Thus, our formulation has four components: the cytosol, the gel, the nuclei, and the activity of PP1 which couples nuclear and cortical dynamics. Since flows in a mid-embryo cross-section have cylindrical symmetry as a first approximation [20], we simplify our modeling framework by developing a 2D description of the flow with the cross-section itself described by an ellipse of major axis 250*μm* and of minor axis 90*μm*. We then obtain the following equations:

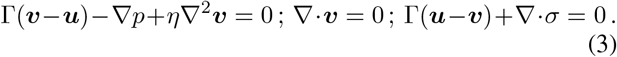

Here, Γ is the gel-sol friction coefficient, ***v*** is the sol velocity, ***u*** is the gel velocity, *p* the pressure field, *η* the shear viscosity of the cytosol and the gel stress tensor *σ* decomposes as :

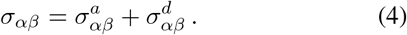

The passive component reads

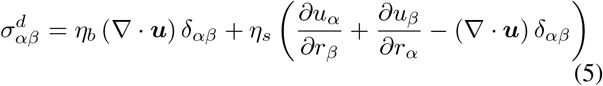

where *η*_*b*_ and *η*_*s*_ are the gel bulk and shear viscosity. The active term models contractions of the actomyosin network driven by gradients in bound myosin concentration. The simplest option for the stress tensor is an isotropic term with saturation :

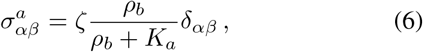

where *ρ*_*b*_ is the concentration of active myosin (myosin bound to actin) [36]. *ζ* is the contraction strength and *K*_*a*_ is the active saturation parameter (see the below section on vorticity for the rationale and tests of this choice).

The cytosol satisfies no-slip boundary conditions **v** | _∂Ω_ = 0 at the boundary of the domain (cortex) ∂Ω. Conversely, the active gel must satisfy the no-penetration condition (**u** · **n**) | _∂Ω_ = 0, where **n** is the normal to the cortex, but can slide along it. The balance of the forces between the cortex and the gel layer in contact with it yields the tangential traction boundary condition (*σ*^*T*^ **n** · **t**) | _∂Ω_ = Ξ. If the gel were free to slip, then Ξ = 0, which is contradicted by the experimental observation of a (weak) backflow following antero-posterior expansions. This observation makes it more appropriate to consider the elastic response discussed hereafter.

As the gel flows, actomyosin filaments attach and detach from the cortex. When detached, filaments are carried by the flow **u**. When attached, filaments are getting strained and, if the contracting force vanishes, they will tend to flow back to their anchoring point. To describe the evolution of the straining displacement *s*, we denote by *k*_*c*_ the binding rate to the cortex and by *τ*_*c*_ the cortical unbinding time. The derivative 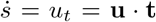 (where **t** is the local tangent to the cortex) if the filament is in the bound state and 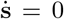 in the unbound state. The probability of the former is 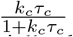 and the duration *t* of the binding event is exponentially distributed as 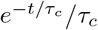. Taking the average to identify typical effects over many filaments, it follows that the average displacement

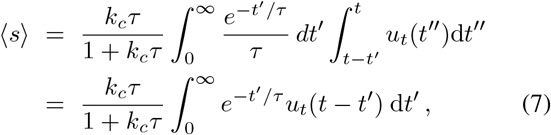

where the second equality is obtained integrating by parts. Given the displacement, the tangential traction is finally :

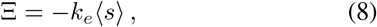

where *k*_*e*_ is the spring constant of the elastic restoring force.

### C. Modeling the control of actomyosin contractility by the cell cycle

Biochemical regulation of myosin activity is controlled by the cell cycle oscillator, viz., the spatiotemporal activity of PP1. Experimental observations suggest the following dependence of PP1 activity on space and time:

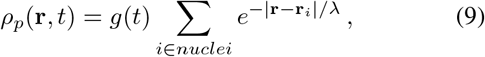

where *g*(*t*) is a suitable oscillatory function that embodies the phase of the cell cycle, **r**_*i*_ is the position of the *i*-th nucleus, **r** is the point of interest, and *λ* ≃ 30*μm* is the decay length of the PP1 activity cloud generated by each nucleus.

We now present the dynamical equations for myosin II and its regulation by the above PP1 field. We consider myosin II to exist in two states: unbound and bound (active) to actin filaments. The unbound and bound states are described by a reaction-diffusion equation:

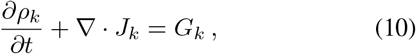

where *J*_*k*_ is the flux of species *k* and *G*_*k*_ is a reaction term. We consider two kind of fluxes: advection and diffusion. The unbound myosin will be advected by the sol, whereas the bound myosin by the gel, so that:

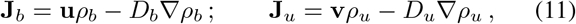

where *D*_*b*_ and *D*_*u*_ are the respective diffusion constants. We consider myosin to diffuse much less when bound than when free in the sol, i.e., *D*_*b*_ ≪ *D*_*u*_.

As for the reaction terms, the two myosin species are coupled by binding/unbinding kinetics, with the total amount of myosin assumed to be constant. Considering these reactions in a linear regime, one obtains:

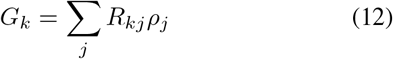

where the elements of the matrix *R* are the reaction rates. Since the total myosin is constant, *R*_*jk*_ = − *R*_*kj*_. We assume that the myosin unbinds at a constant rate so that *R*_*bu*_ = − *R*_*ub*_ = − *k*_*u*_. The activation of myosin is promoted by Rho activity, which is in turn regulated by PP1 and cortical mechanisms. It is observed experimentally that the timing of myosin activation is delayed with respect to that of PP1 and Rho activation [20]. That motivated us to introduce an effective intermediate field *ρ* that responds to PP1 activity with a characteristic time *τ* and mediates the activation of myosin according to the equation :

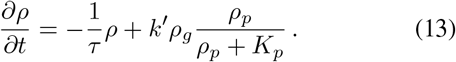

The field *ρ*_*g*_ effectively accounts for the preferential activation of myosin close to the cortex, likely due to the localization of *ρ*-GEF molecules mediating the Rho/myosin activation process, as well as molecules that help organize the cortical actin network. The field rapidly decays away from the cortex as *ρ*_*g*_ = *e*^−*r/μ*^, where *r* indicates the distance from the cortex and *μ* is the characteristic decay length (the value used in the rest of the paper is *μ* = 8*μm*). The field *ρ* controls the rate of myosin activation as :

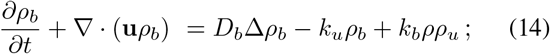

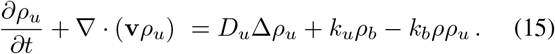

### D. Modeling the transport of nuclei, their divisions and positioning

The last step to complete our model is to define the dynamics of nuclei. Nuclei are advected by the local sol velocity **v** according to the following overdamped equation for the position **r**_*i*_ of the *i*-th nucleus:

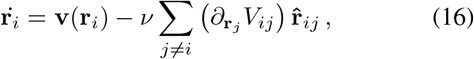

where *ν* is the mobility. The interaction potential *V*_*ij*_ between nuclei *i* and *j* follows an inverse power law with a cutoff :

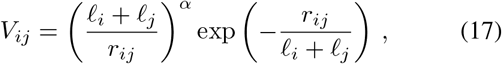

which provides an effective description of the repulsive forces among the two nuclei that result from the action of microtubule spindles, asters and/or actin caps [37, 38]. The simple idea is that when the typical extension of those microtubule structures is comparable/longer than the distance *r*_*ij*_ between the two nuclei *i* and *j*, a repulsive force will result. The length *ℓ*_*i*_ of the microtubule structure radiating from the *i*-th nucleus depends on the phase of its cell cycle. Specifically, based on the observation that microtubule asters are inhibited by Cdk1 and grow at mitotic exit/early interphase [39, 40], we take the following linear dependence on the local PP1 concentration:

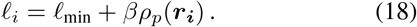

Finally, in (17) we take the exponent *α* = 6, as in molecular dynamics (other choices leave our results unchanged).

Nuclear divisions are regulated by the temporal function *g*(*t*) that controls the PP1 concentration in (9). In particular, the minima of this function mark each mitotic entry, at which point every nucleus is replaced by two nuclei separated by a distance of four microns, centered around the original one. The division axis is randomly chosen for each division event; tests were also performed using a deterministic rule, namely splitting nuclei along the direction perpendicular to the net force exerted on each nuclei. Our main results and conclusions were found unaltered. As the microtubules grow according to (18), the nuclei are pushed apart as a result of the effective potential, thus completing mitosis.

### E. Our model reproduces the large-scale geometry of the embryonic flows

Our model for cytoplasmic flow is defined by the ensemble of Eqs. (3-18). We simulated the model starting with a single nucleus, which undergoes seven cycles of divisions and transport by cytoplasmic flows generated by the above dynamics. Figure 1 shows that the model captures experimental observations. In particular, the large-scale structure of the flow and its timing with the cells cycles are correctly reproduced. The large-scale pattern of the flow features four vortices and a stagnation point that arises near the center of the nuclear cloud where cortical flows converge. A strong antero-posterior (AP) extensive flow is observed during interphase, at the peak of myosin activation and contraction (see Figs. 1A-B). The basic mechanism is as in Ref. [20]: PP1 activity drives cortical myosin gradients, which lead to contractility and motion of the actomyosin gel ; The sol is then entrained by the friction with the gel and its incompressibility produces its ingression in the bulk of the embryo and the four vortices pattern. As for the backflows seen in Figs. 1C-D, they are generated by the elastic restoring force (8). The force pushes back the actomyosin filaments when the action of myosin contractions vanishes as the cell progresses into mitosis. As in the case of the forward flow, the sol is then entrained by friction and creates its own flow. Its structure is similar to the reverse of the forward flow, albeit the amplitude is smaller since the elastic force is weaker than myosin contractility.

**Figure 1.**
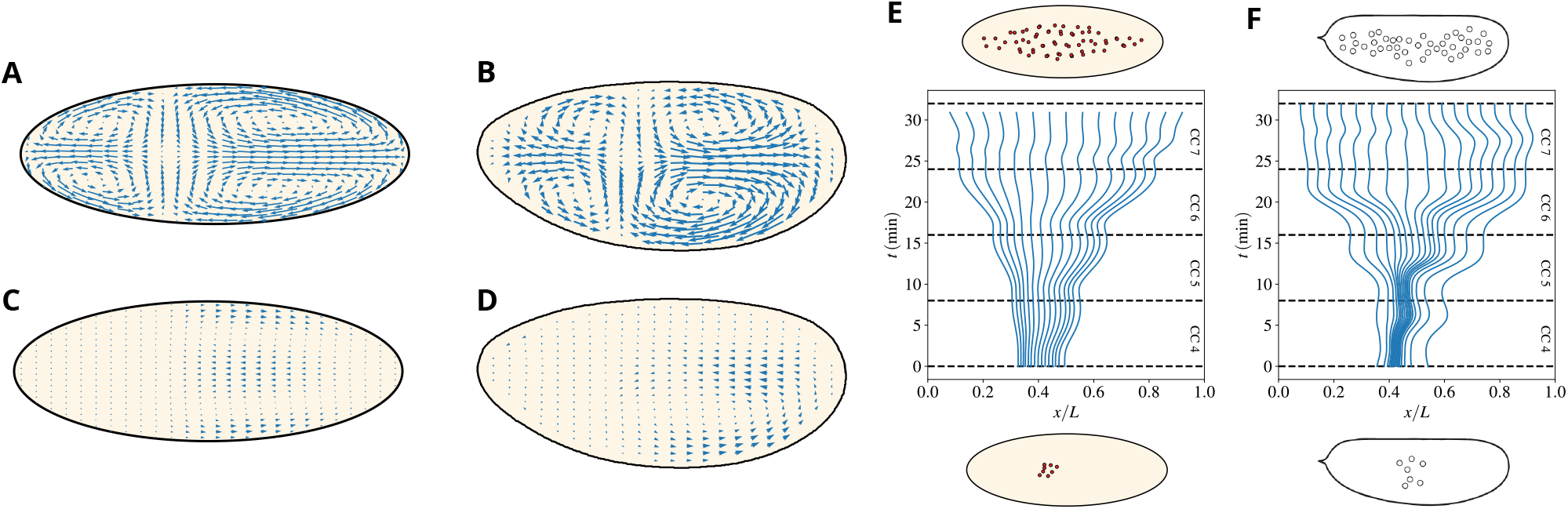
The two-fluid model captures the large-scale feature of embryonic flows. A)-B) Typical cytoplasmic flow observed during the AP expansion phase in our model (A) and experiments (B). C)-D) Typical cytoplasmic backflow observed during the AP contraction phase in our model (C) and experiments (D). The length of the arrows is in the same units as in panel (A,B) so as to highlight the reduced speed of backflow. E)-F) Reconstructed initial distributions to achieve a uniform nuclear distribution at the end of cell cycle 7 in our model (E) and wild-type experiments (F). Particles are uniformly distributed along the AP axis at the end of cycle 7 and simulated (E) or measured (F) cytoplasmic flows are used to evolve their position backward in time until the beginning of cycle 4.

Note that the nuclear positions and flows are strongly coupled by the PP1 profiles being centered around the nuclei. In particular, the positions of the stagnation and the ingression points will move together with the center of the nuclear cloud. This is the basis of the self-correcting properties of cytoplasmic flows that will be discussed later below. As for the dependence on the cycles, cortical and cytoplasmic speeds increase gradually in magnitude from cycles 4–6 and then significantly reduce by cycle 7. The reason for the increase is the growing number of nuclei and the strength of their effects. However, by cycle 7, a new effect sets in : nuclei are almost uniformly spread along the AP axis, which implies that gradients of myosin activation tend to fade and eventually vanish. That is the reason for the minor role of cytoplasmic flows in late cycles.

In sum, Eqs.(3-18) produce effective dynamics to spread uniformly the nuclei over the embryo, as visually conveyed by Fig. 1C. A striking way to condense this message is provided by Figs. 1E-F, which trace back nuclei distributed uniformly at the end of cycle 7. To this aim, we computationally positioned uniformly-distributed particles in the mid-embryo along the AP axis at the end of cycle 7. We then used the simulated or experimentally-measured cytoplasmic flows to evolve their position backward in time until the beginning of cycle 4. The figure clearly shows that nuclei would start from a small cloud centered in the mid-embryo at the beginning of cycle 4 both in the model and the experiments.

### F. Our model captures the myosin dynamics

Experimental data [20] show the peculiar time profiles of Myosin II (see the Supplementary Fig. S3 of Ref. [20] for similar plots for F-actin) for cell cycles 5-6 at the embryo surface and varying distances from it. Peaks of myosin concentration are strongest at the cortex and decay over a few microns away from it, as shown in Fig. 2. The other characteristic feature is that the peaks are delayed at various depths, with the earliest peak at the cortex and the deeper ones progressively delayed.

**Figure 2.**
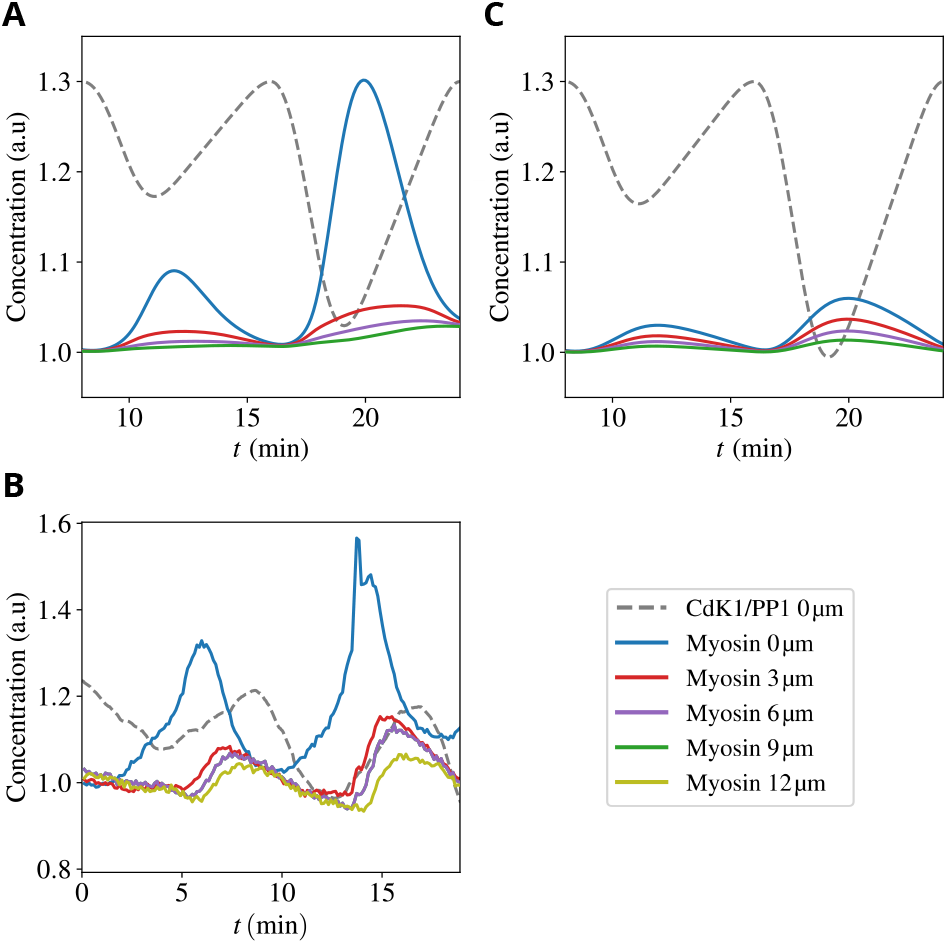
Myosin dynamics in the early embryo. Upper and middle panels: Total concentration of myosin for cell cycles 5-6 at embryo surface (blue line) and varying distances from the surface (see leg-end) in our model (A) and experiments (B). Dotted black line: The Cdk1 to PP1 ratio, which constitutes a proxy for the phase of the cell cycle. Lower panel: Total concentration of myosin for the same conditions as in panel (A) but for a passive gel, i.e., suppressing the active component *σ*_*a*_ of the stress in (4).

The above behavior is well captured by our model, as shown in Figure 2A. The qualitative reason underlying these trends is intuitively understood from the interplay between gel flow and diffusion of unbound components. In a nutshell, the dynamics has two phases. In the first phase, during early interphase, gel flows drive the rapid recruitment of myosin to the cortex where it accumulates in the bound state. The width of its peak close to the cortex reflects the rapid decay of the *ρ*_*g*_ field defined in (13). In the subsequent second phase, at mitotic entry, the PP1 activity decreases, and gel flows dampen. Then, the dynamics is dominated by diffusion of the unbounded components which tend to move from the cortex (zone of high concentration) toward the interior of the embryo. The delays as a function of the depth are caused by the time taken by the excess myosin that unbinds at the cortex to progressively diffuse back. Fig. 2C highlights the role played by the active nature of the gel fluid : if we suppress activity by putting *σ*_*a*_ = 0 in (4), variations in myosin levels are much reduced and the order of the delays among the peaks at different depths is lost.

### G. Our model explains experimental deviations from Stokes’ flow of the vorticity field

The experimental vorticity field in Ref. [20] highlighted the fundamental discrepancy between cytoplasmic and Stokes’ flow explained in the above Section. Conversely, we show here that our model reproduces the presence of extrema of the vorticity field inside the embryo.

By taking the curl of (3), we obtain

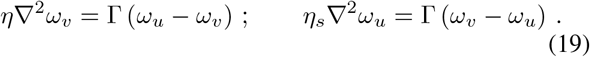

Remark indeed that all the terms in the stress tensor that are ∝ *δ* do not contribute as they yield terms ∝ *ϵ*_*ij*_ Δ_*i*_ Δ_*j*_, where the tensor *ϵ*_12_ = − *ϵ*_21_ = 1 is anti-symmetric. It follows from (19) that, while the sum of the two vorticities *η*_*s*_*ω*_*u*_ + *ηω*_*v*_ is still harmonic, each individual component is not. We conclude that extrema are *a priori* possible for the individual components whilst they are not for their weighted sum. This property is explicitly confirmed in Figure 3.

**Figure 3.**
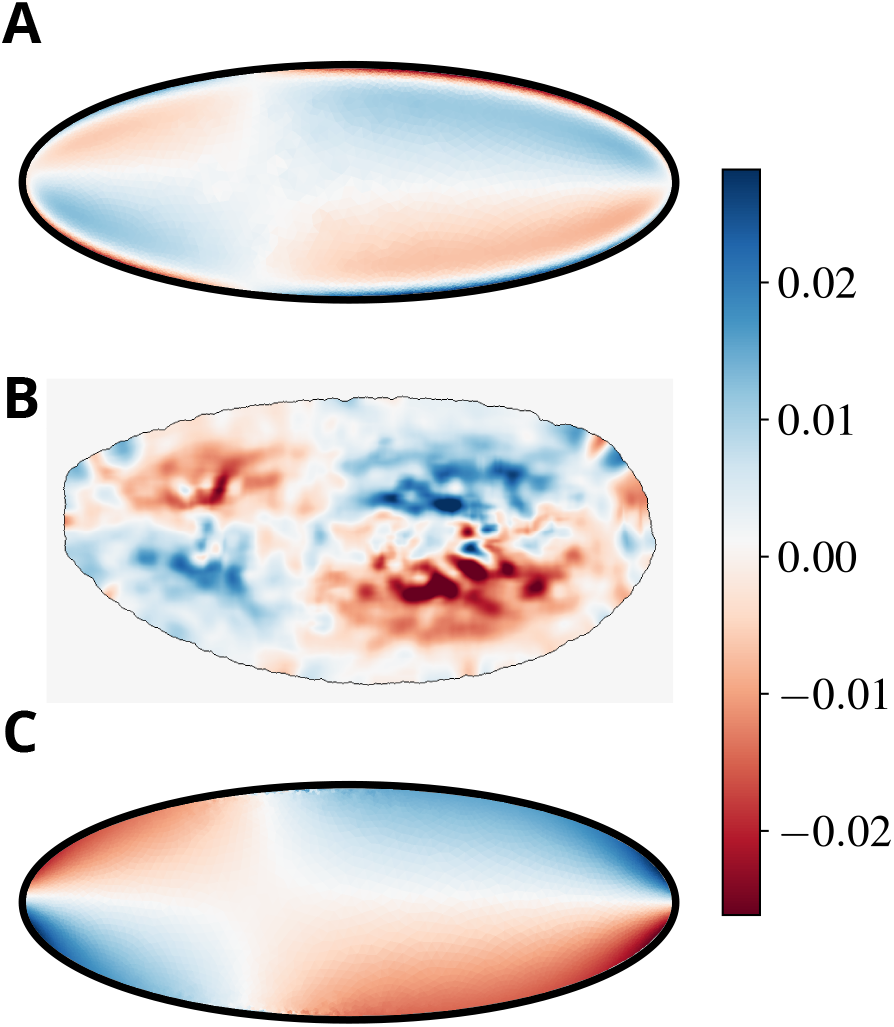
Our model explains experimental observations of the sol vorticity. A)-B) A heatmap showing the vorticity field (*ω*_*v*_ = ∇ × **v**) of the sol flow in our model (A) and in experiments (B). C) The total vorticity *η*_*s*_*ω*_*u*_ + *ηω*_*v*_ (normalized by *η* + *η*_*s*_ to preserve its physical dimensions), showing its extrema at the boundary of the domain, which reflect the harmonic nature of the field in our specific model (see discussion in the body of the paper).

Note that while the dissipative part of the gel stress tensor (5) is fixed, the active component (6) could be non-isotropic and therefore take a non-diagonal form, e.g., in the presence of nematic order [41]. This would *a priori* lead to a non-vanishing contribution to the vorticity balance, which would break the harmonicity of the total vorticity ∝ *η*_*s*_*ω*_*u*_ + *ηω*_*v*_. In the absence of data on the gel velocity, we cannot locate the extrema of the total vorticity and test its harmonicity. Furthermore, we show here that the simple isotropic form (6) captures the main phenomenology available at this stage. That was the rationale for our minimalistic choice of the isotropic form (6), which should of course be tested and possibly revisited as gel flow data become available.

### H. Our model predicts different spatio-temporal flow patterns for gel and cytosol and a sharp micron-size boundary layer at the cortex

An essential feature of our model is the explicit modeling of two fluids : cytosol and gel. Such a model accommodates of course the possibility of an effective single fluid, i.e., that the friction between the two fluids makes their flow similar to each other. The purpose of this Section is to show that that is not the case and the dynamics is genuinely multiphase. This conclusion is evidenced by Figure 4, where snapshots of sol and gel flows (at the same time) are compared. The most striking difference is that the sol flows in its circulating four-vortices patterns with major components along the anteroposterior axis whilst the gel mainly flows from the inside of the embryo towards the cortex. The latter drives the accumulation of gel components to the surface that was quantified in Figure 2 and the corresponding Section. As already mentioned, experimental data on gel flow are not available yet and would be crucial to test our predictions.

**Figure 4.**
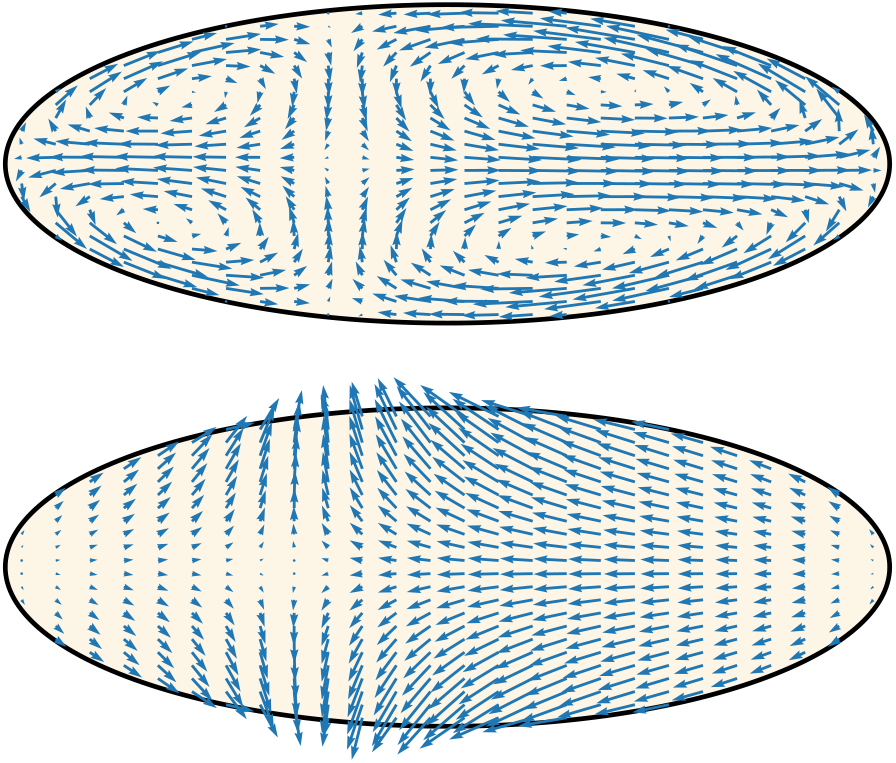
The multiphase nature of the dynamics in our model is germane. The two plots show a typical flow of the sol (upper) and the gel (lower). Note that, contrary to the sol, the gel mainly flows from the inside of the embryo towards the cortex, driving the peaks in myosin concentration in Figure 2.

The difference in the flow patterns of the two fluids should not mislead the conclusion that friction is irrelevant and does not play any role. On the contrary, friction is what drives the flow of the sol, which would be at rest without the entrainment by the gel. Most of the entrainment is provided in the micron-size boundary layer observed in Fig. 5 close to the cortex. The thin, micron-size width of the boundary layer is consistent with the experimental observation that cytoplasmic flows are observed very close to the cortex in spite of the membrane appearing to be essentially immobile and the no-slip boundary condition applying to the sol. The physical explanation and prediction of our model are that a very thin boundary layer exists, where the sol velocity drops abruptly from its bulk value to zero. This would also be consistent with the fact that the typical size of the cortex is of the order of a few hundred nanometers.

**Figure 5.**
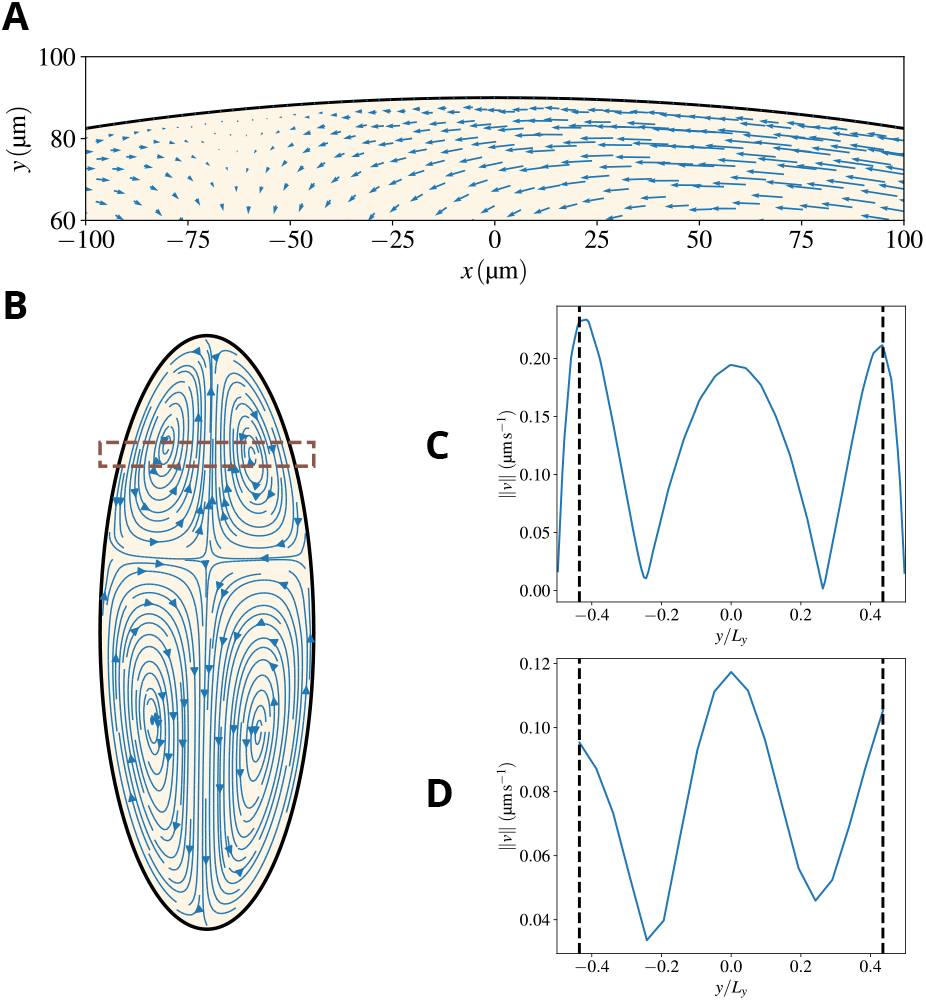
Our model predicts a thin boundary layer close to the cortex. A) A zoom of the region close to the cortex, meant to highlight that substantial cytoplasmic flows are observed relatively close to the cortex, as observed in experiments. In fact, the no-slip boundary condition forces the sol velocity to drop to zero at the cortex but the decrease is sharp and happens in a boundary layer that is micron-thick. This is visually demonstrated by taking the velocity in the box shown in panel (B) and plotting its amplitude *vs* the position (normalized by the width of the embryo at that AP position). Panel (C) shows results for our simulations, and panel (D) for an analogous region in the experiments. The segmented vertical lines represent the point from which no experimental velocities can be resolved, which reinforces the impossibility of resolving a boundary layer such as the one present in the simulations with the available experimental data.

To estimate the width of the boundary layer, we take the difference in (19) :

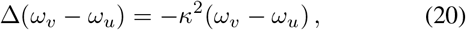

which identifies the reciprocal of a lengthscale 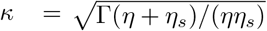 that vanishes in the absence of friction. The parameters that allowed us to reproduce experimental observations yield 1*/κ* ≃ 7*μm*, consistent with previous remarks on the width of the boundary layer. As noted above and shown in Fig. 5D, this fine resolution was not accessible in the experiments in Ref. [20] and constitutes a prediction that awaits experimental validation. From a numerical standpoint, properly capturing the boundary layer required an adaptive mesh, sub-micrometer sized near the embryo boundary, which allowed us to resolve the flow within the boundary layer (see Methods for details).

Note finally that our model provides insights into the physical properties of the actomyosin gel and the sol. In particular, reproducing experimental speeds requires the viscosity of the gel to be a few orders of magnitude larger than the viscosity of the sol. This is consistent with independent experimental data reporting *η*_*gel*_*/η*_*water*_ ≈ 10^5^ [42].

### I. The AP expansion is a self-correcting mechanism for positioning nuclei

The position of the very first nucleus in the embryo is not fixed and subject to fluctuations [20]. The nucleus will then divide rapidly and the resulting cloud of nuclei at the beginning of cycle 4, when the flows start, will reflect its initial position. What is the influence of that position on the structure of the flows and will the final distribution of nuclei, at the end of cycle 7, be affected? In other words, are the flows able to buffer the unavoidable shifts and distortions of the nuclear cloud at the beginning of cycle 4 and still achieve a uniform spreading of the nuclei?

The answer to this question is summarized by Fig. 6, where we have simulated various initial positions and reported the difference of the nuclear positions with respect to the reference configuration where the initial nucleus is placed at 40% of the total AP length of the embryo. Several remarkable features can be noted. First, irrespective of the initial position, the cloud of nuclei converges to very similar final configurations. Second, the role of forward and backward flows is apparent from the figure: the distance to the reference configuration decreases during the forward phases and flattens or even increases during the phases of backflow. Third, the final differences between the actual and the reference positions are only a few microns, defining the degree of robustness of the process.

**Figure 6.**
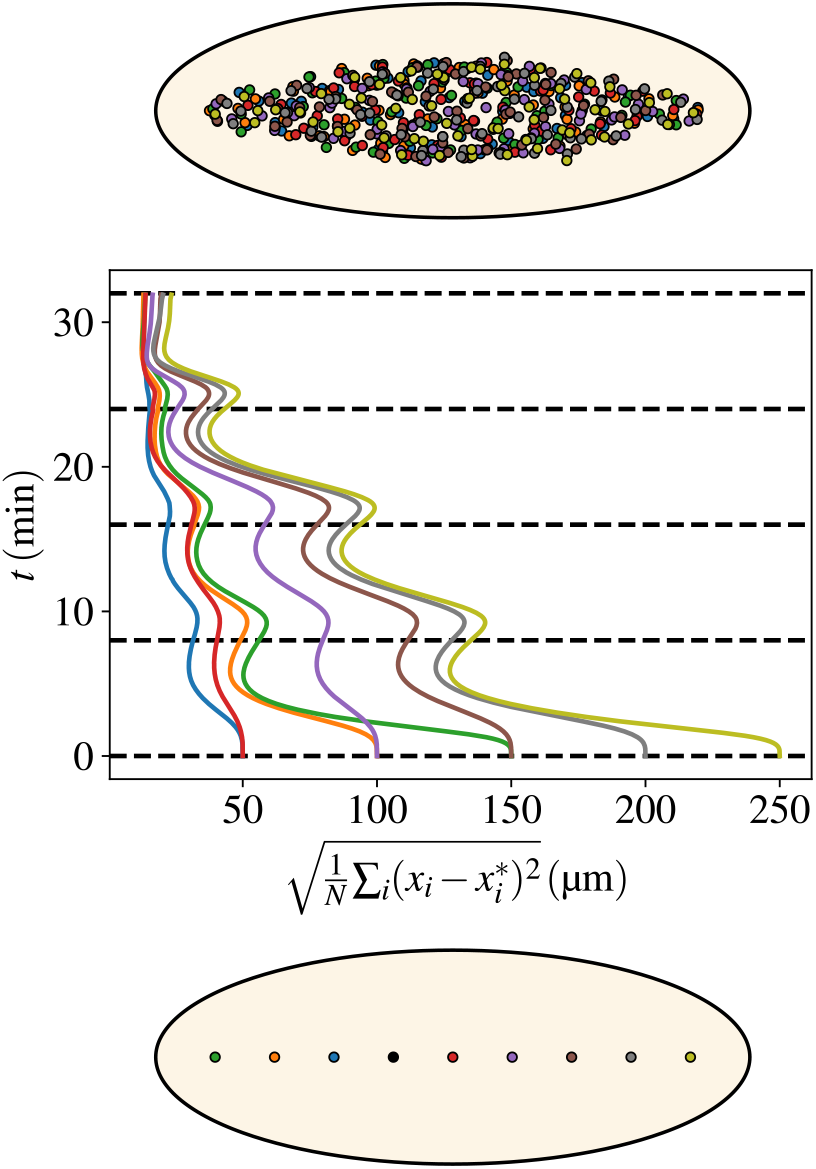
Flows ensure a uniform distribution of nuclei along the AP embryonic axis irrespective of the initial nucleus’ location. Bottom: A series of positions (coded by different colors) for the first nucleus that starts the division cycles. The nine different locations go from 10% to 90% of the embryo AP length. The reference configuration has the nucleus placed at 40% of the embryo AP length. Middle: The evolution in time (flowing upwards) of the distance with respect to the reference configuration. The distance is defined as 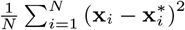, where *N* is the total number of nuclei (at that time), 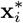 and **x**_*i*_ are the positions of nuclei in the reference or the displaced configurations. The best matching that minimizes the total distance between the two sets of nuclei is obtained by using the Belief Propagation algorithm described in the Methods. Colors of the curves correspond to the initial positions in the bottom panel. Three pairs correspond to the two sides of the reference positions and the last two curves refer to the initial positions at 80% and 90% of the AP length. Top: The final configurations of nuclei (for the whole ensemble of colors). Note that all colors are mixed up, witnessing the self-correcting nature of the AP spreading process. That is shown more quantitatively by the middle curves, which all reduce to values corresponding to distances of a few microns distance between pairs of nuclei of the various configurations.

The reasons underlying this striking property of the cytoplasmic flow reside in what was stressed in previous sections: myosin accumulation and contractions are localized at the position of the nuclear cloud. That defines the location where the sol will flow in the bulk and extend. If the initial location is sufficiently central, as it typically is, then four vortices are created and an extension both on the anterior and posterior sides is produced. Conversely, if the initial position is very anterior (or posterior), then the flow points in the opposite direction toward the posterior (or anterior). The net effect is that the modified dynamics compensates for the displacement of the initial nucleus and readjusts the final position of the nuclei as demonstrated by Figure 6.

We also tested the ability of the flows to center nuclei in response to displacements in directions perpendicular to the AP axis (see Fig. S1). We found that the flows are able to center the nuclei and ensure that the nuclear cloud sits in the middle of the embryo. The reason for this is intuitive. If nuclei are closer to one side of the cortex than the other, PP1 activity and myosin recruitment are higher on that side and, as a consequence, that side will experience a stronger contraction and flow, which will help to center the nuclei.

### J. The impact of cytoplasmic flows on the formation of the Bicoid morphogen gradient

The cytoplasmic flows at cell cycles 4-6 are concomitant with the initial stage of the establishment of the Bicoid gradient [43]. Bicoid is a morphogen that generates an exponential gradient essential to pattern the AP axis of the embryo [44, 45]. The formation of the gradient is controlled by the localization of *bcd* mRNA to the anterior of the embryo [46]. The mRNA is translationally silent until fertilization when the localized production coupled with protein motion starts the process that eventually establishes the morphogenetic gradient [43, 46]. The movement of the protein has been hypothesized to be dominated by diffusion [47]. However, formation of the Bicoid gradient is not fully understood and it has been suggested that flows might in fact play a role in ensuring that the Bicoid gradient achieves the appropriate length [48]. However, this hypothesis has remained largely untested due to the lack of quantitative information on the structure of cytoplasmic flows.

To test the influence of cytoplasmic flows on the formation of Bicoid gradients, we used the flow generated by the model described above and simulated the dynamics of Bicoid transport by using measured mRNA distributions and assuming that protein production begins at fertilization. Bicoid molecules are dispersed by the joint effect of molecular diffusion and cytoplasmic flows. The minimal assumption is that Bicoid initially does not preferentially localize to the cortex or the bulk. Results for the evolution of the Bicoid concentration are reported in Fig. 7. A color-coded profile of Bicoid at various times is shown in panel A. The profiles of Bicoid concentration at cycles 4-7 reported in panel B are in agreement with the experimental curves reported in [49]. In our model, we can easily turn off the flows and ascertain their effect on the gradients’ formation. The corresponding curves illustrate that the differences with and without flows are minor. More specifically, in the anterior part of the embryo, the flow at the cortex tends to push Bicoid toward the mid-embryo and the curves with the flows on are therefore slightly higher than without. The effect is reversed beyond mid-embryo but remains minor. Moreover, effects on Bicoid spreading are predicted to be temporary: by cell cycles 7-8, Bicoid distribution is essentially indistinguishable with and without flows. Additional evidence is provided by Fig. S2. We conclude that flows have a minor influence on the formation of the Bicoid gradient.

**Figure 7.**
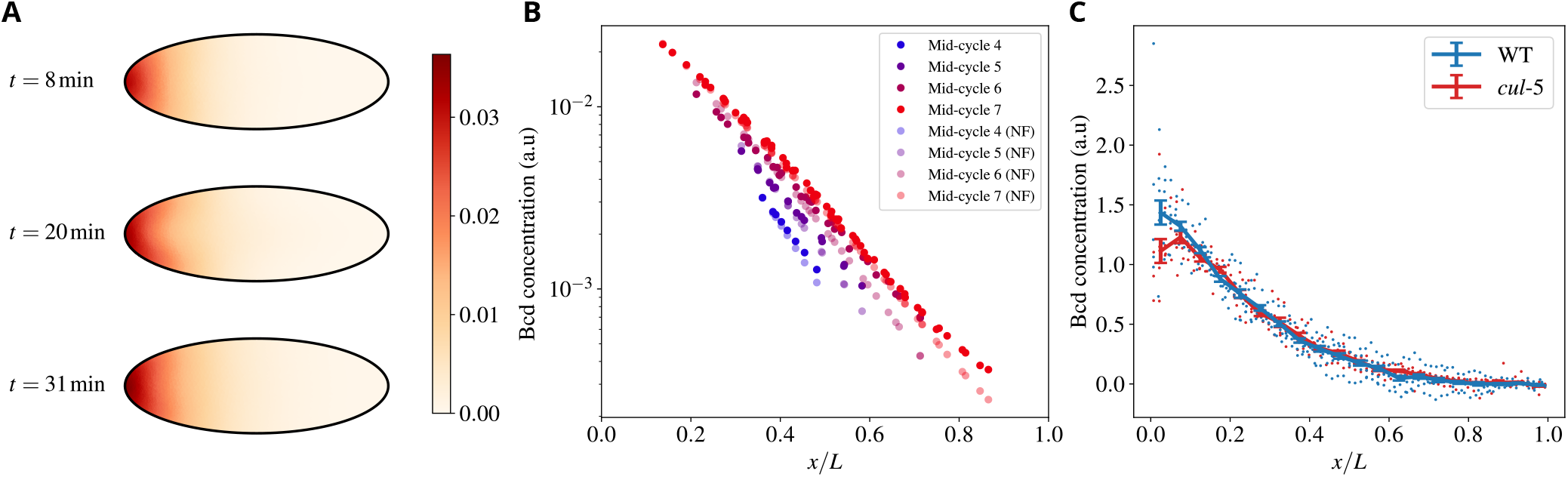
Embryonic cytoplasmic flows weakly affect the establishment of the Bicoid morphogenetic gradient. A) Heatmap showing the Bicoid concentration profile in the embryo, and the characteristic half-moon shape with the concentration higher at the cortex than in the bulk. B) The concentration of Bicoid *vs* the (normalized) position along the AP axis for various cycles, as indicated in the color legend. NF stands for “No Flow”, i.e., situations where cytoplasmic flows were suppressed. Values are reported at the position of each nucleus. C) Experimental comparison of the Bicoid gradient at cell cycle 13 in wild-type and *cullin-5* mutant embryos, where flows are strongly suppressed [29]. Datapoints were binned in each case, with the continuous line and error bars representing each bin’s average and standard error respectively.

To test this prediction experimentally, we compared the Bicoid gradient in wild-type embryos and mutant embryos, featuring severely reduced cytoplasmic flows. To this end, we used *cullin-5* (*shackleton*) mutant embryos, which we have previously shown to display strongly reduced cortical contractions and cytoplasmic flows [29]. Consistent with a minor and transient role for flows in controlling the Bicoid gradient, we found that the gradient at cell cycle 13 is essentially indistinguishable in wild-type and *cullin-5* mutant embryos.

In sum, even though the presence of cytoplasmic flows might *a priori* influence the formation of the Bicoid gradient, in practice the structure of the flows actually observed in the embryo is such that their influence is minor and they weakly and transiently affect the formation of Bicoid gradients.

### K. Our model predicts the effects of embryo geometry on cytoplasmic flows that are experimentally verified

As a further test of our model, we performed various simulations altering the geometry of the embryo. The resulting dynamics is well captured by the following intuitive considerations, which run similar to those explaining the self-correcting nature of the AP expansion demonstrated in Figure 6.

The cortex points which are closest to the nuclear cloud define the regions where myosin-II accumulates and they potentially determine the locations of the strongest contraction. In the wild-type and the usual geometry, these points will determine where the gel flows are directed and define the corners of the vortices observed in the sol flows. An important role is played by the elliptic geometry of the embryo since the transversal distance between the AP axis and the cortex reduces as one moves toward the poles. A first expected effect is that, as the ratio between the major and minor axes of the ellipse reduces to unity, the dynamics is going to become more isotropic, and therefore the ratio between longitudinal and transverse speeds should tend to unity. The second expected effect is a global reduction in the overall speed, which can be linked to two different causes. First, due to changes in embryo geometry on average nuclei will be further away from the cortex, and the PP1 cloud will not activate as much myosin as in the wild-type case. Furthermore, as the egg becomes more symmetric, the shape of the PP1 isolines becomes closer to the shape of the egg, thus reducing the myosin gradients and the strength of the flows.

To test our predictions, we generated embryos of different geometry by using a knockdown of Fat2 (Fat2 RNAi), a major regulator of the elongation of the egg chamber, in somatic cells of the female ovary [50, 51]. Notably, these experiments are performed by directing the expression of the RNAi transgene to the somatic cells of the egg chamber, thus embryos of different geometry but genetically wild-type can be generated, see Figure 8A. These embryos show patterns of flow that differ significantly as compared to the wild-type. In particular, the ratio between longitudinal and transverse speeds, as well as the overall speed, reduce (see panels B and C) in agreement with our theoretical predictions.

**Figure 8.**
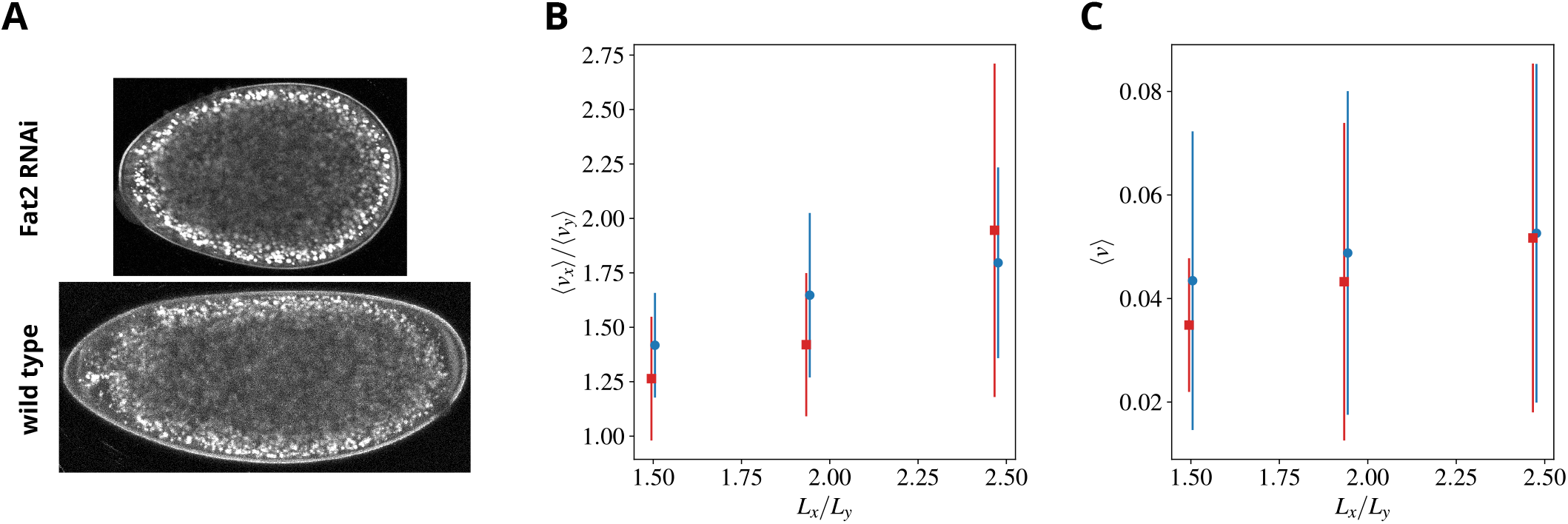
Predictions and experimental verification of flows in rounded embryos. A) A wild-type and a round embryo are shown in the lower/upper panels, respectively. Mutants are generated by using a knockdown of Fat2 (Fat2 RNAi), a major regulator of the elongation of the egg chamber. Panels B) and C) show the ratio between the embryo-averaged longitudinal and transverse sol speeds and the embryo-averaged sol speed, respectively, time-averaged over CC6. Simulation/experimental data are shown in blue circles/red squares, respectively. Error bars represent the standard deviation of the space averages during CC6.

## II. DISCUSSION

We have presented a theoretical framework that quantitatively captures the main mechano-chemical couplings underlying the cytoplasmic flows observed in the early *Drosophila* embryo. As in the great majority of developmental and cellular conditions, Reynolds numbers in the fly embryos are low, and fluids are well described by neglecting accelerations (Stokes’ conditions). The behavior of a Stokes fluid is determined by its velocity at the boundary of the region of interest [35], a property that underlies most of previous works on cytoplasmic flow. There, the velocity at the boundary of the cell or the embryo was prescribed, e.g., by the measured velocity, and it was then shown that the cytoplasmic flow is well described by solving Stokes’ equations with the prescribed boundary conditions [16, 19, 30]. The velocity at the boundary often involves active mechanisms generated, for instance, by active actomyosin contractions. In multicellular systems, active myosin at the cell-cell interfaces can drive tissue-wide cellular flow, as shown in *Drosophila* gastrulation [52]. Global morphogenetic flow is accurately predicted by the spatial distribution of myosin motors. Similarly, a tensile ring in the quail embryo drives movement in the gastrulating amniote embryo [53, 54]. Here, we have explicitly coupled the mechanics of Stokes flows to the cell cycle via some of its major chemical components (Cdk1 and PP1). Nuclei in the fly syncytium regulate the balance between Cdk1 and PP1, which itself controls the activation of myosin via Rho. We have shown that fundamental and empirical reasons impose a multiphase model that features a mixture of a passive and an active fluid. The active gel can slide along the cortex and generates the cortical forces that propel the ensemble of the two fluids and the transport of nuclei in the embryo. The passive phase satisfies no-slip boundary conditions and the sol fluid is thus at rest at the plasma membrane, which is adjacent to the immobile *Drosophila* vitelline membrane. A major result emerging from our work is that the two phases move very differently, and a single-fluid description is insufficient to capture the complexity of cytoplasmic flows in fly embryos. Previous discrepancies from a Stokes’ flow of the sol vorticity [20] can thus be understood as reflecting the transfer of momentum between the active and passive phases due to their mutual friction. The entrainment of the passive fluid by its active counterpart concentrates in a thin boundary layer close to the cortex, which constitutes another striking result emerging from our work. In the boundary layer, the velocities of the two phases change sharply to accommodate their respective boundary conditions. The corresponding thickness is predicted to be a few microns.

Our framework recapitulates previous observations and, most importantly, offers new predictions to inspire experiments. The results on round embryos that we presented here provide a notable example of theoretical predictions that we could already confirm experimentally. Numerical simulations of our model predict the extent to which the amplitude and geometry of cytoplasmic flows would change with embryo shape. These predictions were supported by experiments in which embryos of different geometry were generated via genetic engineering. In particular, by knocking down Fat2 expression (via RNAi) in the somatic cells of the egg chamber, we could obtain embryos having a round geometry [50, 51]. In these embryos, the observed cytoplasmic flows were found to be in agreement with our theoretical predictions. Another example of what theory can uniquely offer is provided by our predictions on the role of cytoplasmic flows in the establishment of Bicoid gradients. Specifically, we capitalized on the possibility of direct and unequivocal comparison of gradients with and without cytoplasmic flows to demonstrate their minor role in the establishment process of Bicoid gradients.

In the future, there are several new experiments suggested by our theoretical results. A first example is provided by the robustness of the spreading process with respect to the position of the initial nucleus that we highlighted here. The procedure that we have implemented in numerical simulations suggests set-ups to reproduce it experimentally. For example, one could use optogenetic control of myosin activity to displace the position of nuclei at early cycles [20] and observe whether or not flows compensate for the displacement and still drive uniform positioning in later cycles.

A crucial test of our theoretical framework will be the experimental verification of the gel flow and the presence of a sharp boundary layer. Experimental challenges will need to be overcome to obtain accurate measurements of the flow of the actomyosin gel. First, it is difficult with current fluorescent probes to distinguish whether myosin is in the active (bound) or inactive (unbound) state. Second, imaging of myosin deep in the embryo does not show significant features to infer possible flows. On a positive note, it is relatively straightforward to measure the total concentration of myosin and quantify myosin concentration at different distances from the cortex. Our simulations recapitulate the measured concentration data (see Figure 2), arguing that we captured the essential features of the gel flow. Our simulations predict that the gel flow drives cortical recruitment of active myosin, which is the main driving force of the nuclear spreading process. Approaches based on Fluorescence Recovery After Photobleaching (FRAP) could be used to infer average flows across large regions of the embryo [20, 55]. FRAP of fluorescently tagged myosin near the cortex could potentially confirm that the gel flow is directed mainly towards the cortex (as we predicted here for the gel) rather than sliding parallel to it (as we predicted for the sol). However, testing this prediction experimentally will be strongly influenced by the ratio of the concentration of active and inactive myosin and the results could be difficult to interpret. Thus, we propose that novel experimental approaches are needed to accurately measure the gel flow. Developing such methods will not only reveal fundamental insights into the mechanisms of cytoplasmic flows but will also allow testing the hypothesis that the viscosity of the two fluids is very different, a property that is essential in our model to generate the observed flows of cytosol. Similarly, it will be important to develop experimental approaches to resolve the thin (few *μm*) boundary layers predicted by our theory. Particle Image Velocimetry allows tracking flows near the cortex [20] but assaying the localization of the tracked features relative to the plasma membrane has not been analyzed carefully to date.

Future experimental observations of the gel flow will be important not only to confirm the current theoretical framework but also to refine and advance it. The isotropic form of the active stress tensor that we have taken here was dictated by the parsimony principle and the fact that no current experimental observation forced us to introduce more sophisticated hypotheses (and the resulting additional fields). It is however quite conceivable that nematic and/or polar effects are in fact present and our description should be upgraded to take them into account. A detailed classification of the additional fields and couplings can be found in Ref. [41]. An experimental observable that will be highly informative in that respect is the gel vorticity, which would allow us to construct the total vorticity. As we have shown here, isotropic forms of the active stress tensor cannot produce vorticity. The total vorticity is then harmonic, which strongly constrains its spatial structure (its extrema are confined to the boundary of the embryonic domain). Polar and nematic effects break isotropy and generically lead to the production of vorticity and the breaking of harmonicity. It would then be important to measure the total vorticity and assess whether or not its spatial structure is consistent with the harmonic property.

In sum, the combination of discriminating experimental tests, new predictions, and suggested novel experiments presented here constitute exemplary instances of the value of the interplay between theory, numerical simulations, and quantitative experiments that we see as the exciting way forward in the field of developmental biology and embryology.

## III. METHODS AND MATERIALS

### A. Fly Stocks

To image and quantify the Bicoid gradient, we used an EGFP-Bcd line (Bloomington stock 29018). For the analysis of the gradient in *cullin-5* embryos, we used the following females : EGFP-Bcd/EGFP-Bcd ; His2Av-mRFP/His2Av-mRFP ; *shkl*^*GM*130^/*shkl*^*GM*163^. To obtain round embryos, we crossed flies carrying a *UAS:Fat2* RNAi transgene (VDRC 5098, kind gift of Sally Horne-Badovinac) to flies expressing Gal4 under the *trafficjam* promoter to restrict expression in the somatic cells of the egg chamber (kind gift of Sally Horne-Badovinac). We then selected female flies carrying both transgenes and set up cages to collect embryos where the activity of Fat2 was reduced specifically in the somatic cells of the female ovary. In some experiments, embryos also expressed PCNA-TagRFP to visualize nuclei.

### B. Embryo Manipulations

Following collection, embryos were dechorionated with 50% bleach for 1 min, rinsed with water, mounted in halocarbon oil on a gas-permeable membrane, and covered with a glass coverslip. To visualize cytoplasmic flows, yolk granules were stained by permeabilizing embryos with a solution of 10% CitraSolv in water for 2 minutes and immersing them in Trypan Blue for 1 minute.

### C. Microscopy

Imaging experiments were performed with an upright Leica SP8 confocal microscope, a 20 X /0.75 numerical aperture oil immersion objective, an argon ion laser, and a 561-nm diode laser, as described in Ref. [20]. For the analysis of the Bicoid gradient, images were acquired on the mid-sagittal plane with a time resolution ≃ 12*s*. For the analysis of cytoplasmic flow in wild-type embryos, we used data in Ref. [20], where we acquired stacks of raw confocal sections (800 × 400 pixels, pixel size: 0.727 *μ*m) of yolk (Trypan Blue) and nuclei (PCNA-TagRFP) with an axial distance of 50-60 *μm* and sampling of 20 s. For the round embryos, we acquired images (1024×1024 pixels, pixel size: 0.568 *μ*m) of yolk at about 40 *μ*m from the cortex and sampling ≃ 10 − 20*s*.

### D. Image and data analysis

#### Particle Image Velocimetry

Raw confocal images of yolk granules were Gaussian-filtered (width of 10 *μ*m and standard deviation 6 *μ*m) to increase the signal-to-noise ratio as a pre-processing image analysis step. Cytoplasmic velocity fields were measured by means of Particle-Image-Velocimetry. Briefly, stripes of 35 *μ*m (Anterior-Posterior direction) by 15 *μ*m (Dorsal-Ventral or lateral direction) were used as templates and probed within regions of 60 *μ*m by 30 *μ*m to find the best correlation spots, with a threshold correlation coefficient of 0.7. PIV was calculated for 10,000 unique points randomly distributed in the embryo at each time interval. A sampling of around 20 to 30 s was used to get reliable local displacements while maintaining high correlations. The obtained velocity fields were time-averaged over a range of 10 s and linearly interpolated on a square grid with 4 *μ*m spacing.

#### Quantification of the Bicoid gradient

The nuclear segmentation masks of the *Drosophila* embryos were generated with Ilastik 1.3.3 software [56] by using the Pixel Classification pipeline. The segmented nuclear region was binned along the AP axis, with bin width 7.27 *μ*m (10 pixels, pixel size = 0.727 *μ*m). The average pixel intensity of the EGFP-Bicoid channel was calculated for each binned region of the nuclear mask. Next, 11 consecutive frames (frame rate = 12.56 s) were manually selected right before the mitosis of cell cycle 13 (cc13). The average of these 11 frames was taken for each bin to generate the Bicoid gradient profile. The gradients were normalized as follows: first, the positions of the bins were normalized by the length of the embryo; second, an offset was determined by the average intensity of the 10% posterior-most bins and subtracted from the profile; third, the value at each bin was normalized by the average intensity (post-offsetting) of the 10-20% anterior-most bins. The final profiles of 6 wild-type embryos and 4 *cullin-5* mutant embryos were reported.

### E. Numerical simulations

#### Space and time discretization

We performed numerical simulations by using a finite-element method, implemented in a custom FreeFem++ code. Different meshes were generated for each geometry with an adaptive edge size, such that the boundary layer could be well resolved. The target minimum edge size was chosen as *h*_*min*_ = 0.75 μm, and we used a fixed timestep Δ*t* = 3 × 10^−2^ s. The element types were chosen as MINI elements (Linear element + order 3 bubble) for ***u*** and ***v*** ; discontinuous linear elements for *ρ*_*b*_ and *ρ*_*u*_ ; linear elements for the remaining fields. The mix of MINI elements for the velocity space and linear elements for the pressure space ensures the solution is unique, satisfying the discrete inf-sup condition [57]. Since the flow and myosin dynamical equations are coupled, we sequentially solved them. Specifically, at timestep *n* we first calculate ***v***^*n*^ and ***u***^*n*^ from 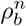 and Ξ^*n*^ ; then we obtain 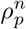 from the nuclei positions ***r***^*n*^, and determine the microtubule lengths *ℓ*^*n*^ ; 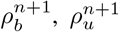 and 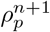 are calculated next, followed by ***r***^*n*+1^ from ***v***^*n*^ and *ℓ*^*n*^ ; finally, we determine ⟨*s*⟩ ^*n*+1^ and Ξ^*n*+1^ from ***u***^*n*^.

#### Stokes equations parameters

The value of the elastic constant that allowed us to reproduce the strength of the backflow was *k*_*e*_ = 6.67 N*/*m^3^. After experimenting in a wide range of values, the contraction strength was set to *ζ* = 1.42 × 10^1^ Pa, and the bound myosin saturation parameter was set to *K*_*a*_ = 1. Similarly, the friction coefficient was set to Γ = 2.2 × 10^7^ Pa*/*m^2^, and *η*_*s*_ = *η*_*b*_ = 1 Pa s. The non-penetration and traction boundary condition for the gel using Nitsche’s method [58], with *β* = 10^7^ as penalization constant.

#### Myosin dynamics

The equations (13,14,15) are conveniently written in a compact form as

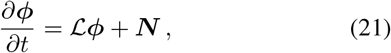

where ***ϕ*** = *ρ* for (13) and ***ϕ*** = (*ρ*_*b*_, *ρ*_*u*_) for Eqs. (14,15). ℒ represents linear parts of the equation, viz. ℒ = −1*/τ* and 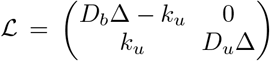, respectively. Conversely, 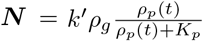 and ***N*** = (*k*_*b*_*ρρ*_*u*_ − ∇ · (***u****ρ*_*b*_), −*k*_*b*_*ρρ*_*u*_ − ∇ · (***v****ρ*_*u*_)) for the two cases. The equation (21) is solved as

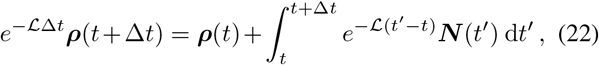

and the integral was approximated by using *N* (*t*′) ≃ *N* (*t*) + (*N* (*t*) − *N* (*t* − Δ*t*))(*t*′ − *t*)*/*Δ*t* to finally obtain :

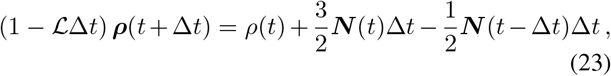

Based on the experimental delay between PP1 and myosin activation peaks, we chose *τ*_*I*_ = 7.5 × 10^−1^ min, *k*′ = 1.6 and *K*_*p*_ = 5 for (13). As for Eqs. (14,15), the selected parameters were *k*_*u*_ = 8.33 × 10^−1^ s^−1^, *k*_*b*_ = 6.67 × 10^−1^ s^−1^, *D*_*u*_ = 1.67 × 10^2^ μm^2^*/*s and *D*_*b*_ = 1.67 μm^2^*/*s The initial conditions were *ρ*_*u*_ = 1 and *ρ*_*b*_ = 0 everywhere. At the boundary, the fields satisfy no-flux boundary conditions, which ensure that the total amount of myosin in the embryo is constant.

#### Bicoid dynamics

We modeled the dynamics by the equation

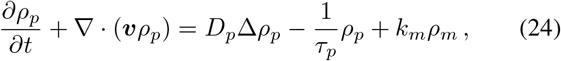

with the diffusivity *D*_*p*_ = 5.0 μm^2^*/*s, the decay time *τ*_*p*_ = 60 min and production rate *k*_*m*_ = 1.67 × 10^−4^ s^−1^. The Bicoid mRNA distribution *ρ*_*m*_ decays exponentially from the anterior pole 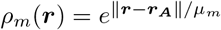, where ***r***_***A***_ is the position of the anterior pole, and *μ*_*m*_ = 25 μm is the decay length, that is ∼ 5% of the embryo length. By defining 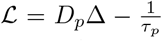, and *N* (*t*) = − ∇ · (***v****ρ*_*p*_) + *k*_*m*_*ρ*_*m*_, we can solve this equation as explained above. Our initial conditions have *ρ*_*p*_ = 0 everywhere.

#### Cortical actomyosin dynamics

As mentioned in the main text, backflows are explained by the elastic recoil of actomyosin filaments, which attach and detach to the cortex. If *i* denotes a generic boundary node and ⟨*s*_*i*_ ⟩ represents the mean displacement of filaments attached at *i* (positive for counter-clockwise displacements), the evolution of the displacement is given by :

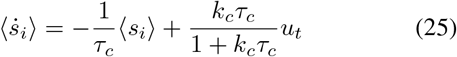

where *u*_*t*_ is the component of the gel velocity tangent to the boundary at the displaced location. This equation was solved as described above with ℒ accounting for the decay term with typical rate 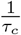, and *N* (*t*) the non-homogeneous last term on the r.h.s of the equation. A value *τ*_*c*_ = 4 min reproduced the backflow delay and duration. Similar values for the cortical elasticity relaxation timescale have been reported in previous experimental works using ferrofluids at the beginning of cellularization [59]. We have used *k*_*c*_ = 4 min^−1^.

#### Nuclear dynamics

Based on the ellipsoidal flow geometry observed in previous experiments [20], we exploited the cylindrical symmetry around the AP axis and performed simulations in a twodimensional (2d) domain representing a mid-embryo slice. In order to take into account the third (*z*) dimension into account, we proceeded as follows. First, from the *x* and *y* positions of nuclei, we built Δ_*x*_ and Δ_*y*_, the maximum nucleus-nucleus distance in *x* and *y*, and their center of mass. Second, a 3d ellipsoid is built at the center of mass and semimajor axes Δ_*x*_*/*2, Δ_*y*_*/*2 and Δ_*y*_*/*2. The last step enforces the cylindrical symmetry. Third, *z*-coordinates of nuclei are assigned so that each one of them lies at the boundary of said ellipsoid. For each nucleus there are two possible *z*-coordinates : if the previous *z*-coordinate is non-zero, its sign is preserved ; else, it is chosen at random. Finally, distances required for microtubule-mediated interactions (17), PP1 activity (9) and pair matching algorithm, all considered 3d distances.

Eqs. (16) for the nuclei were integrated using a 1-step Euler method. To obtain nuclear separations comparable to the experiments, we chose *ℓ*_*min*_ = 1 μm, *β* = 1 μm. At mitotic exit, we duplicated each nucleus, choosing a random axis of division and placing two daughter nuclei at a distance *r*_*mit*_ = 2 μm along the division axis. Initial conditions in the unperturbed embryo considered a single nucleus along the AP axis and 50 μm to the left of the center.

#### Pair matching

To produce Fig. 6, we had to find the best matching between two sets of particles (Nuclei). Matching two sets of particles in two different image frames is conveniently formulated as the matching in a bipartite graph so that a certain edge-weight dependent quantity is minimized [60]. This problem is well-known in graph theory [61], and efficient algorithms are available [62]. In our case, for each timestep, the two sets {***r***_***i***_(*t*)} and 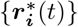 represent the set of nuclei positions in a reference simulation and another simulation where the initial nucleus has been displaced. To find the matching which minimizes the RMS particle-particle distance, we implemented the simplified min-sum algorithm described in [62], with a weight function 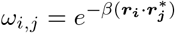, and *β* = 0.002.

## IV. ACKNOWLEDGMENTS

We thank the Bloomington Stock Center and S. Horne-Badovinac for providing stocks. We thank A. Chao, V. Deneke and L. Hayden for experimental support, help with data analysis and discussion. We thank all the members of the Di Talia and Vergassola labs for discussions. We thank P. Adhyapok, O. Afonso, A. Box, A. De Simone, M. Kramar and C. Lang for initial observations on Bicoid gradients in wild type and *cullin-5* mutant embryos, made as a part of the Summer Research Course at the Kavli Institute of Theoretical Physics (KITP), UCSB. Our visit to KITP was supported by NSF grant no. PHY-1748958, NIH grant no. R25GM067110, and the Gordon and Betty Moore Foundation grant no. 2919.01. This work was partially supported by the NIH: R01-GM136763 (to S.D. and M.V) and by “Associazione Italiana Ricerca sul Cancro” MFAG-2020 n. 25040 (to A.P.). C.H. was supported by a PhD grant from ED564 ‘Physique en Ile de France’.

