## Supplementary Information for "Two-fluid dynamics and micron-thin boundary layers shape cytoplasmic flows in early *Drosophila* embryos"

### Notation

- $\Omega$: embryo domain.
- $\partial\Omega$: embryo boundary.
- $\hat{\boldsymbol{n}}$ and $\hat{\boldsymbol{t}}$: normal and tangent vectors to the embryo boundary.
- $\boldsymbol{w}^v$, $\boldsymbol{w}^u$, $\boldsymbol{w}^p$, $\pi^b$, $\pi^u$ and $\pi^p$: basis functions for the $\boldsymbol{v}$, $\boldsymbol{u}$,$p$, $\rho_b$, $\rho_u$ and $\rho_p$ spaces respectively.
- $\boldsymbol{A} : \boldsymbol{B} = \sum_{ij} A_{ij} B_{ij}$

When the corresponding finite element space is discontinuous, additional quantities must be defined. Let $F$ be the shared edge between the adjacent triangles $T_1$ and $T_2$. Then:

- $h_F$: length of $F$.
- $\hat{\boldsymbol{n}}_F$: normal vector to $F$.
- $[[g]] = g(T_1) - g(T_2)$: jump discontinuity of $g$ across $F$.
- $\{\{g\}\} = \frac{1}{2}\left(g(T_1) + g(T_2)\right)$: average of $g$ across $F$.

### Weak formulation of the flow equations

After multiplying Eqs. 3 by their respective basis functions, and integrating over the whole embryo, we get the following equations to solve by our finite element scheme:

$$\eta \int_\Omega (\nabla \boldsymbol{v}) : (\nabla \boldsymbol{w}^v) - \int_\Omega p(\nabla \cdot \boldsymbol{w}^v) + \Gamma \int_\Omega (\boldsymbol{v} - \boldsymbol{u}) \cdot \boldsymbol{w}^v = 0 \tag{S1}$$

$$\eta_s \int_\Omega (\nabla \boldsymbol{u}) : (\nabla \boldsymbol{w}^u) + \mathcal{C}(\eta_s, \eta_b, \boldsymbol{u}, \boldsymbol{w}^u) + \mathcal{D}(\gamma_n, \boldsymbol{u}, \boldsymbol{w}^u) + \Gamma \int_\Omega (\boldsymbol{u} - \boldsymbol{v}) \cdot \boldsymbol{w}^u = -\zeta \int_\Omega \frac{\rho_b}{K_a + \rho_b} (\nabla \cdot \boldsymbol{w}^u) + \int_{\partial\Omega} \Xi(\boldsymbol{w}^u \cdot \hat{\boldsymbol{t}}) \tag{S2}$$

$$\int_\Omega w^p(\nabla \cdot \boldsymbol{v}) - \gamma_p \int_\Omega p w^p = 0 \tag{S3}$$

where:

$$\mathcal{C}(\eta_s, \eta_b, \boldsymbol{u}, \boldsymbol{w}^u) = \eta_b \int_\Omega (\nabla \cdot \boldsymbol{u})(\nabla \cdot \boldsymbol{w}^u) + \eta_s \int_\Omega \left( \frac{\partial u_y}{\partial x}\frac{\partial w^u_x}{\partial y} + \frac{\partial u_x}{\partial y}\frac{\partial w^u_y}{\partial x} - \frac{\partial u_y}{\partial y}\frac{\partial w^u_x}{\partial x} - \frac{\partial u_x}{\partial x}\frac{\partial w^u_y}{\partial y} \right) \tag{S4}$$

arises from the gel compressibility, and:

$$\mathcal{D}(\gamma_n, \boldsymbol{u}, \boldsymbol{w}^u) = \gamma_n \int_{\partial\Omega} (\boldsymbol{u} \cdot \hat{\boldsymbol{n}})(\boldsymbol{w}^u \cdot \hat{\boldsymbol{n}}) - \int_{\partial\Omega} (\hat{\boldsymbol{n}}^T \boldsymbol{\sigma}^d[\boldsymbol{u}]\hat{\boldsymbol{n}})(\boldsymbol{w}^u \cdot \hat{\boldsymbol{n}}) - \int_{\partial\Omega} (\hat{\boldsymbol{n}}^T \boldsymbol{\sigma}^d[\boldsymbol{w}^u]\hat{\boldsymbol{n}})(\boldsymbol{u} \cdot \hat{\boldsymbol{n}}) \tag{S5}$$

is an extra term that enforces the no-penetration boundary condition via penalization (Symmetric Nitsche's method). We chose $\gamma_n = 10^7$. $\gamma_p = 10^{-10}$ is a small term that ensures the algorithm converges to a well-defined pressure.

### Weak formulation of the myosin continuity equations

$$\left(\frac{1}{\Delta t} + k_u\right) \int_\Omega \rho_b^{n+1}\pi^b + \mathcal{A}(D_b, \gamma_g^b, \rho_b^{n+1}, \pi^b) = \frac{1}{\Delta t}\int_\Omega \rho_b^n \pi^b + k_b \int_\Omega \rho^n \rho_u^n \pi^b + \frac{3}{2}\mathcal{B}(\rho_b^n, \boldsymbol{u}^n, \pi^b) - \frac{1}{2}\mathcal{B}(\rho_b^{n-1}, \boldsymbol{u}^{n-1}, \pi^b) \tag{S6}$$

$$\frac{1}{\Delta t}\int_\Omega \rho_u^{n+1}\pi^u - k_u \int_\Omega \rho_b^{n+1}\pi^u + \mathcal{A}(D_u, \gamma_g^u, \rho_u^{n+1}, \pi^u) = \frac{1}{\Delta t}\int_\Omega \rho_u^n \pi^u - k_b \int_\Omega \rho^n \rho_u^n \pi^u + \frac{3}{2}\mathcal{B}(\rho_u^n, \boldsymbol{v}^n, \pi^u) - \frac{1}{2}\mathcal{B}(\rho_u^{n-1}, \boldsymbol{v}^{n-1}, \pi^u) \tag{S7}$$

where:

$$\mathcal{A}(D,\gamma,\rho,\pi) = D\int_\Omega (\nabla\rho)\cdot(\nabla\pi) + \gamma D\sum_F \frac{1}{h_F}\int_F [[\rho]][[\pi]] - D\sum_F\int_F (\{\{\nabla\rho\}\}\cdot\boldsymbol{n}_F)[[\pi]] - D\sum_F\int_F [[\rho]](\{\{\nabla\pi\}\}\cdot\boldsymbol{n}_F) \quad \text{(S8)}$$

is the diffusion term, including discontinuity penalization, and:

$$\mathcal{B}(\rho,\boldsymbol{u},\pi) = -\int_\omega \nabla\cdot(\rho\boldsymbol{u})\pi + \sum_F\int_F (\boldsymbol{u}\cdot\hat{\boldsymbol{n}}_F)[[\rho]]\{\{\pi\}\} - \frac{1}{2}\sum_F\int_F |\boldsymbol{u}\cdot\hat{\boldsymbol{n}}_F|[[\rho]][[\pi]] \quad \text{(S9)}$$

is the advection term, including penalization and upwinding.

We chose $\gamma_g^b = 5\times 10^2$ and $\gamma_g^u = 5\times 10^3$ as the discontinuous jump penalty coefficients for $\rho_b$ and $\rho_u$ respectively.

After calculating $\rho_b^{n+1}$ and $\rho_b^{n+1}$, a 2-D limiter (modified *minmod*) is applied as described in [1]. The parameters chosen in our case were $M = 5\times 10^{-1}$ for the limiter curvature cutoff, and $\theta = 1$ as its comparison parameter.

**Weak formulation of the Bicoid continuity equation**

$$\left(\frac{1}{\Delta t}+\frac{1}{\tau}\right)\int_\Omega \rho_p^{n+1}\pi^p + \mathcal{A}(D_p,\gamma_g^p,\rho_p^{n+1},\pi^p) = \frac{1}{\Delta t}\int_\Omega \rho_p^n\pi^p + \mathcal{B}(\rho_p^n,\boldsymbol{v}^n,\pi^p) + k_m\int_\Omega \rho_m^n\pi^p \quad \text{(S10)}$$

where we chose $\gamma_g^p = 5\times 10^3$.

**A simplified model for the actomyosin concentration dynamics**

The same qualitative features as the myosin dynamics in Fig. 2 are obtained in the following simplified one-dimensional model. A constant unit concentration of scalar $\theta_u$ is present in the negative half-line in the unbound $u$ form. Particles are reflected at the origin (meant to represent the cortex), i.e., reflecting boundary conditions are imposed there. At the origin, the scalar can interconvert between the $u$ and the $b$ forms at rates $k\theta_u(0)$ and $\theta_b/\tau$.

During a first phase of accumulation at the cortex, a constant velocity $V$ is present in the positive direction, which dominates diffusion and replenishes the amount that is converted into the $b$ form at the origin. The solution in this first phase is calculated easily, unit for all the positions but the origin. The equations for the values at the origin (unbound and bound, respectively) are

$$\dot{\theta}_0 = -k\theta_0 + V + \frac{\phi_0}{\tau}\,; \qquad \dot{\phi}_0 = k\theta_0 - \frac{\phi_0}{\tau}\,, \quad \text{(S11)}$$

where we have already used that $\theta_{-1} = 1$. With $V = 0$, the values settle to $\theta_0 = 1$ and $\phi_0 = k\tau$, which are approached exponentially with rate $(1+k\tau)/\tau$. In the presence of $V$, there is a pile-up and both values go up in time as $Vt/(1+k\tau)$ and $Vtk\tau/(1+k\tau)$.

The above process keeps going on until the cell cycle gets to its end, when the flow $V$ and the attachment rate $k$ vanish. During this second phase of movement, the bound scalar at the origin is released and diffuses back toward the bulk. The re-injection rate is $ke^{-t/\tau} = \phi_0(t)/\tau$, where $t$ is counted from the beginning of this second phase. The time-profile at a distance $x$ from the cortex is then proportional to $e^{-t/\tau}\int_0^t G(x,s)e^{s/\tau}\,ds$ where $G(x,s)$ is the diffusive propagator at distance $x$ and time $s$ with reflecting boundary conditions at the origin. The latter is calculated by the method of images [2], and, since the source is located close to the reflecting boundary, $G(x,s)$ is essentially the diffusive propagator if $x$ is not small. We can finally calculate the above time-profile and check that it has a structure similar to Fig. 2 with peaks that are shifted later and later in time as the distance to the cortex increases.

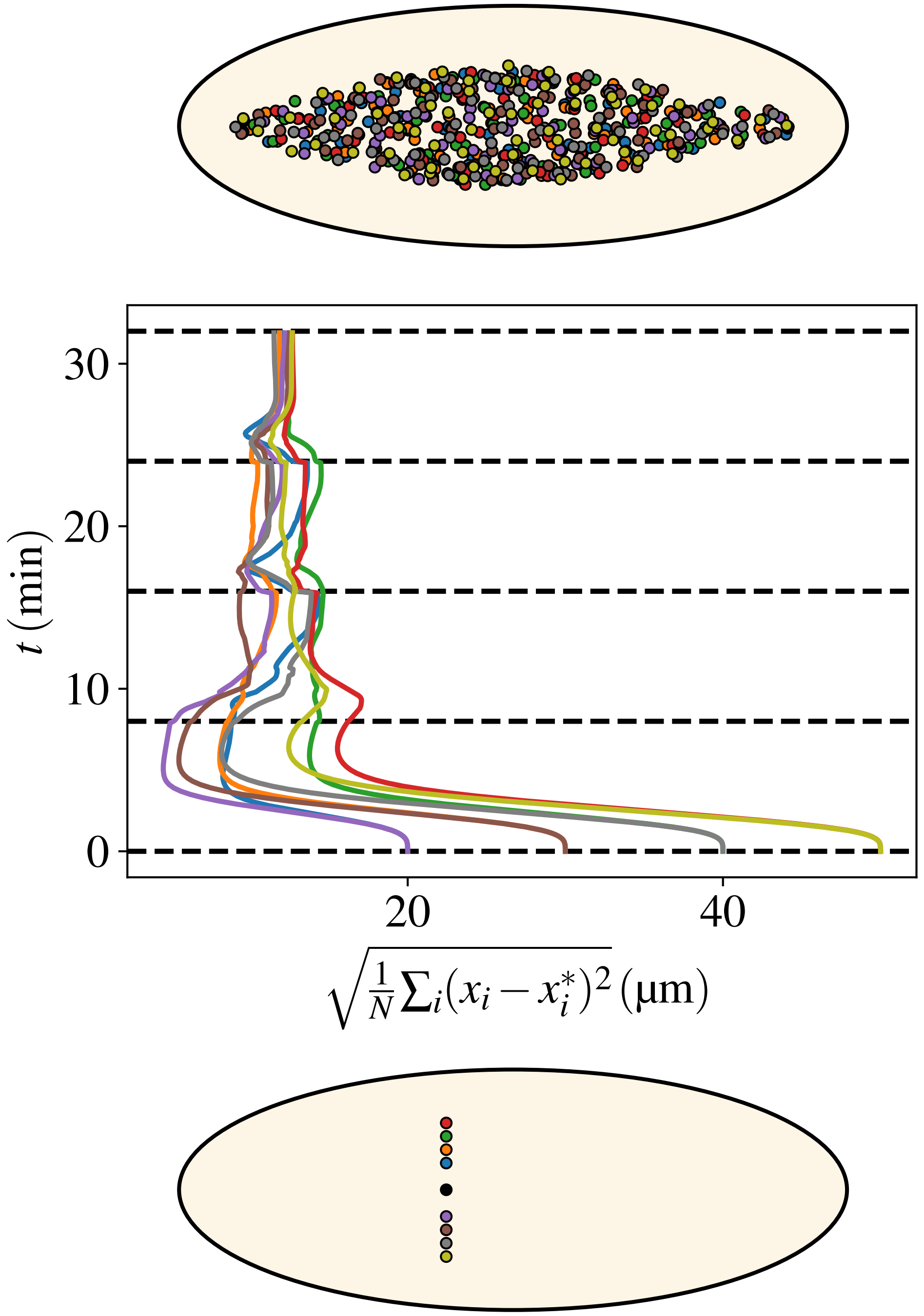


Figure S1. **Flows ensure a uniform distribution of nuclei along the AP embryonic axis irrespective of the initial nucleus' distance to it.** Bottom: A series of positions (coded by different colors) for the first nucleus that starts the division cycles. The nine different locations consider the reference configuration at the embryo AP axis, and four pairs displaced 20 µmto 50 µm away from said axis. Middle: The evolution in time (flowing upwards) of the distance with respect to the reference configuration. Distance is defined and calculated as in Fig. 6. Colors of the curves correspond to the initial positions in the bottom panel. Top: The final configurations of nuclei (for the whole ensemble of colors). Note that all colors are mixed up, witnessing the self-correcting nature of the AP spreading process. That is shown more quantitatively by the middle curves, which all reduce to values corresponding to distances of a few microns distance between pairs of nuclei of the various configurations.

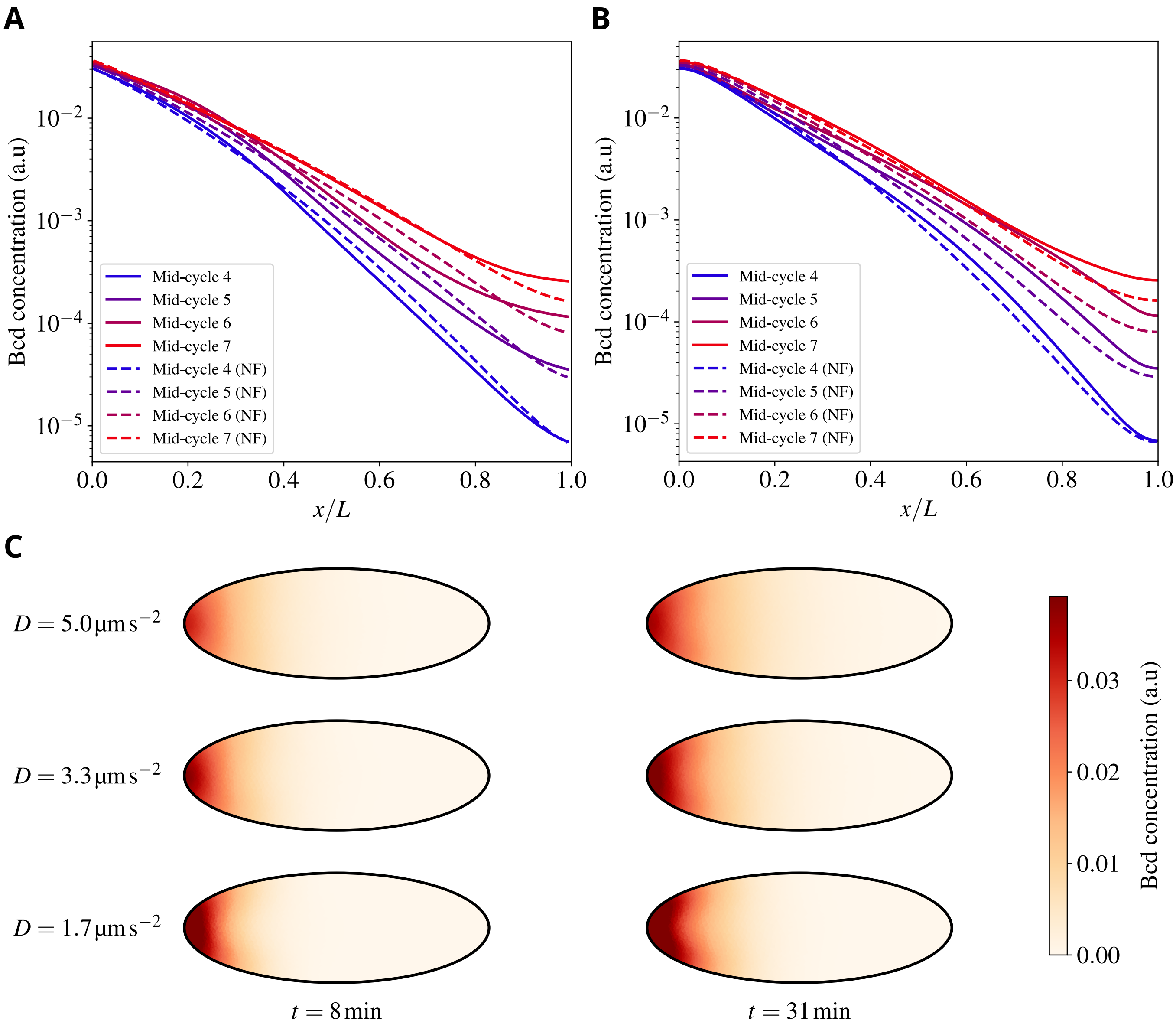


Figure S2. **Embryonic cytoplasmic flows weakly affect the establishment of the Bicoid morphogenetic gradient.** A)-B) Mid-embryo (A) and cortical (B) Bicoid concentrations *vs* the (normalized) position along the AP axis for various cycles, as indicated in the color legend. NF stands for "No Flow", i.e., situations where cytoplasmic flows were suppressed. C) Heatmaps showing the Bicoid concentration field at two different times after fertilization and three different Bicoid diffusivities.

**Movie S1.** Nuclear spreading: with flows (Top) and without flows (Bottom)

**Movie S2.** Self-correcting positioning: four initial starting positions.

**Movie S3.** Chemical species dynamics: PP1 (Top left), total myosin (Top right), bound myosin (Bottom left) and unbound myosin (Bottom right).

[1] D. Di Pietro and A. Ern, *Mathematical Aspects of Discontinuous Galerkin Methods*, Vol. 69 (2012).
[2] S. Redner, *A Guide to First-Passage Processes* (Cambridge University Press, Cambridge, 2001).
